# Processing cookies formulated with goat cream enriched with conjugated linoleic acid

**DOI:** 10.1101/542977

**Authors:** Ana C.S. Costa, Diego E. Pereira, Caio M. Veríssimo, Marcos A. D. Bomfim, Rita C.R.E. Queiroga, Marta S. Madruga, Susana Alves, Rui J.B. Bessa, Maria E.G. Oliveira, Juliana K.B. Soares

## Abstract

Goat fat is one of the most important sources of conjugated linoleic acid (CLA), a fatty acid which has health benefits. However, CLA consumption is limited to meats and milk products as CLA is generated in ruminants. This study aimed to replace vegetable fat by goat cream enriched with CLA. Four cookie recipes were developed with only the fat source being different: CVF – vegetable fat; CB – butter; CGC – goat cream without CLA; CGCLA – goat cream with CLA. Cookies were evaluated according to physical (color and texture) and physical-chemical parameters (lipids, proteins, total sugars, fiber, ash, moisture and Aw), Consumer Testing (n = 123) and lipid profile. The CGCLA presented higher values in the color parameters, and the higher and the lower scores in relation to hardness were 5.54 (CB) and 2.21 (CVF), respectively. Lipids and total sugars varied inversely, and the highest percentages of lipids were in the CVF and CG samples, which obtained lower total sugar content. There was no difference in the acceptance and preference of the four formulations, and the formulations with the goat creams (CG and CGCLA) were as accepted as CFV. The lipid profile of the cookies presented CFV with the highest percentage of trans fatty acids (TFA) with 16.76 %. CGCLA presented 70 % more CLA in relation to CB and CGC, thus certifying that CLA was present in relevant quantities in the CGCLA, even after cooking. The CGCLA is a biscuit with higher levels of CLA, and in this study it was possible to verify that the goat milk cream enriched with CLA can be used in producing cookies which adds functional and nutritional properties to them and offers other alternatives to produce food from goat’s milk cream.

## Introduction

The trans-industrial fatty acids (TFA) present in hydrogenated vegetable fat are associated with adverse health impacts, particularly on changes in the lipoprotein profile in blood plasma which are directly related to cardiovascular diseases and other non-chronic diseases communicable diseases (CNCD). Trans-rumen fatty acids (TFAr) are the TFA formed by the biohydrogenation process of linoleic and linolenic acids in ruminants, and by the enzymatic action of Δ 9 -dessaturase on vaccinic acid such as conjugated linoleic acid (CLA) [1]. CLA is scientifically known as having functional anti-carcinogenic, anti-obese, abdominal fat reducing, anti-atherogenic and immunomodulatory properties [2, 3].

Due to TFAr production occurring in ruminants, CLA is characteristic of these animals and is found in the lipids fraction of its milk or meat. Caprine fat is one of the main sources of CLA, in addition to having other physiological benefits such as lowering cholesterol levels without altering triglycerides, HDL, TGO or TGP levels, inducing neurodevelopment stimulus, reducing anxiety and reducing intestinal inflammation in rats [4-6].

Hydrogenated vegetable fat is used industrially in most bakery and confectionery products such as biscuits and cookies, and the use of this fat is associated with the physicochemical and technological properties of food such as texture, plasticity and high melting point [7, 8]. Due to these characteristics of industrial trans fats, it is a challenge to develop products such as biscuits, and especially those rich in fat as cookies are, using other lipid sources. Cookies are biscuits that have high levels of sugar and fat and low levels of water (1-5%), and can be elaborated from different flavors [9, 10].

In this context, the objective of this work was to develop cookie type biscuits elaborated from the caprine cream enriched with CLA, aiming to obtain an alternative matrix for the consumption of this TFAr which is only found in dairy and meat products.

## Material and methods

### Material

The samples of goat’s milk cream were supplied by the Brazilian Agricultural Research Company of goats and sheep, Sobral - CE. The production of enriched CLA cream was enabled by manipulating the diet of the animals with the addition of soybean oil by EMBRAPA [11].The other ingredients used in the cookie formulations were purchased from markets in Joao Pessoa-PB and Recife-PE.

### Lipid profile of fats

Samples of fats (VF, vegetable fat, B, butter, G, goat cream, and GCLA, goat cream enriched with CLA) were analyzed for their lipid profiles. The procedure of transesterification of the samples was performed according to Sukhija and Palmquist (1988) and Palmquist and Jenkins (2003), with adaptations [12. 13]. The fatty acids were analyzed according to methyl esters by gas chromatography with flame ionization detection (GC-FID), using a QP2010-plus Shimadzu chromatograph (Shimadzu, Kyoto, Japan) equipped with a silica-fused (SP-2560, 100 m × 0.25 mm × 0.20 μm, Supelco, Bellefonte, PA, USA).

During the analysis, the injector and detector were maintained at 250°C and 280°C, respectively. Helium was used as the entrainment gas at a constant flow rate of 1 ml/min and 1 μl of sample was injected. The oven temperature was programmed to start at 50°C, then increased by 50°C every minute until it reached 150°C, remaining for 20 minutes, next increased by 1°C every minute until it reached 190°C for 1 minute, then increased 2°C every minute until reaching 220°C, and finally remaining at 220°C for 30 minutes. The identified fatty acids were express as percentages and are shown in Table 1.

**Table 1.** Fatty acid profile present in fats in percentage.

| **FATTY ACIDS** | **VF<sup>a</sup>** | **B<sup>b</sup>** | **G<sup>c</sup>** | **NCLA<sup>d</sup>** |
| --- | --- | --- | --- | --- |
| **SATURATED** |  |  |  |  |
| C4:0 | - | 3,32 | 2,29 | 2,61 |
| C6:0 | - | 2,06 | 2,75 | 2,27 |
| C8:0 | 0,01 | 1,19 | 3,27 | 1,84 |
| C10:0 | 0,01 | 2,58 | 11,14 | 5,28 |
| C12:0 | 0,07 | 2,98 | 4,37 | 1,95 |
| C14:0 | 0,13 | 10,91 | 10,15 | 6,14 |
| C15:0 | 0,02 | 1,92 | 1,38 | 1,01 |
| C16:0 | 12,77 | 31,01 | 25,55 | 21,01 |
| C17:0 | 0,03 | 1,43 | 1,09 | 0,97 |
| C18:0 | 11,20 | 11,32 | 12,13 | 21,20 |
| C20:0 | 0,39 | 0,15 | 0,29 | 0,37 |
| C21:0 | 0,03 | 0,03 | 0,04 | 0,04 |
| C22:0 | 0,44 | 0,06 | 0,07 | 0,11 |
| C23:0 | 0,05 | 0,03 | 0,02 | 0,02 |
| C24:0 | 0,16 | 0,05 | 0,03 | 0,02 |
| **TOTAL SFA** | **25,32** | **69,01** | **75,41** | **64,81** |
| **MONOUNSATURATED** |  |  |  |  |
| C14:1C9 | - | 0,94 | 0,11 | 0,05 |
| C16:1C7 | 0,01 | 0,23 | 0,27 | 0,29 |
| C16:1C9 | 0,06 | 1,38 | 0,41 | 0,31 |
| C17:1C9 | 0,01 | 0,26 | 0,16 | 0,10 |
| C18:1T6+T8 | 3,98 | 0,26 | 0,16 | 0,52 |
| C18:1T9 | 3,04 | 0,19 | 0,14 | 0,46 |
| C18:1T10 | 7,90 | 0,26 | 0,16 | 0,45 |
| C18:1T11 | 4,81 | 1,90 | 0,65 | 4,45 |
| C18:1T12 | 3,31 | 0,26 | 0,17 | 0,60 |
| C18:1C9 | 30,86 | 21,41 | 19,21 | 22,79 |
| C18:1T15 | 1,60 | 0,18 | 0,10 | 0,35 |
| C18:1C11 | 2,25 | 0,44 | 0,30 | 0,34 |
| C18:1C12 | 8,90 | 0,13 | 0,09 | 0,24 |
| C18:1C13 | 0,46 | 0,06 | 0,02 | 0,05 |
| C18:1T16+C14 | 0,47 | 0,30 | 0,18 | 0,48 |
| C18:1C15 | 0,44 | 0,10 | 0,03 | 0,09 |
| C20:1 | 0,16 | 0,04 | 0,04 | 0,06 |
| TOTAL MUFA | 67,80 | 28,27 | 22,15 | 31,57 |
| **POLYUNSATURATED** |  |  |  |  |
| C18:2N6 | 6,22 | 1,16 | 1,56 | 1,52 |
| C18:3N-6 | - | 0,02 | 0,02 | 0,01 |
| C18:3N3 | 0,20 | 0,40 | 0,22 | 0,14 |
| **CLAC9T11** | - | **0,87** | **0,34** | **1,69** |
| C20:3N-6 | - | 0,04 | 0,02 | 0,01 |
| C20:4N-6 | - | 0,08 | 0,16 | 0,13 |
| C20:5N-3 | - | 0,03 | 0,02 | 0,01 |
| C22:5N-3 | - | 0,06 | 0,05 | 0,04 |
| C22:6N-3 | - | 0,01 | 0,02 | 0,03 |
| TOTAL PUFA | 6,42 | 2,66 | 2,43 | 3,58 |
| **TRANS** | 25,11 | 3,34 | 1,54 | 7,31 |

<sup>a</sup> VF: vegetable fat; <sup>b</sup> B: butter, <sup>c</sup> G: goat cream; <sup>d</sup> GCLA: goat cream with CLA, expressed as a percentage.

### Cookie production

Four formulations of cookies were developed by only varying the fat source between each of them, being CVF - vegetal fat; CB - butter; CG - goat cream without enrichment of CLA; CGCLA - goat cream with CLA enrichment. Preliminary tests were carried out until the ideal formulations were determined before the final composition of the cookies. The inputs and quantities used are describe in Table 2. After preparation, the cookies were baked for 15 minutes at 165°C in a conventional oven.

**Table 2.** Ingredients used in the formulations of cookies for 100 g of fresh dough.

| Ingredients (g) | Cookies |  |  |  |
| --- | --- | --- | --- | --- |
|  | CVF<sup>a</sup> | CB<sup>b</sup> | CG<sup>c</sup> | CGCLA<sup>d</sup> |
| Maize starch | 17,84 | 17,84 | 17,84 | 17,84 |
| Rice flour | 11,74 | 11,74 | 11,74 | 11,74 |
| Vegetable fat | 18,78 | - | - | - |
| Butter | - | 18,78 | - | - |
| Goat cream<br>whitout<br>enrichment of<br>CLA | - | - | 18,78 | - |
| Goat cream<br>with enrichment<br>of CLA | - | - | - | 18,78 |
| Brown sugar<br>mascavo | 18,78 | 18,78 | 18,78 | 18,78 |
| Chestnut | 11,74 | 11,75 | 11,74 | 11,74 |
| Raisin | 9,4 | 9,4 | 9,4 | 9,4 |
| Egg | 9,4 | 9,4 | 9,4 | 9,4 |
| Xanthan gum | 0,47 | 0,47 | 0,47 | 0,47 |
| Cinnamon powder | 0,24 | 0,24 | 0,24 | 0,24 |
| Clove powder | 0,24 | 0,24 | 0,24 | 0,24 |
| Yeast chemical | 0,94 | 0,94 | 0,94 | 0,94 |
| Salt | 0,47 | 0,47 | 0,47 | 0,47 |

<sup>a</sup> CVF - vegetable fat; <sup>b</sup> CB - butter; <sup>c</sup> CG - goat cream; <sup>d</sup> CGCLA - goat cream with CLA.

### Physical analyses

#### Instrumental color

Determination of the cookies’ instrumental color was performed in a CR-300 Minolta colorimeter (MinoltaCo., Osaka, Japan) according to the CIE-lab system [14]. In the CIELAB colorimetric space defined by L*, a*, b*, the L* coordinate corresponds to the brightness, ranging from 0 (black) to 100 (white), while a* and b* refer to the green (-) chromaticity coordinates/red (+) and blue (-)/yellow (+), respectively. The measurements were performed with the apparatus previously calibrated in the reflectance modality with specular reflection excluded and using reference plates. A colon at the top of the cookies was for the measurement and then a dot at the bottom.

#### Instrumental texture

Determination of the cookies’ texture was performed in a TA XT Plus texturometer from Extralab Brazil. The data obtained were analyzed by StableMicroSystems software/TE32L/Version 6.1.4.0, England. Each sample was placed horizontally on the platform and cut in half with a blade probe (HDP/3PB), pre-test speed of 1 mm/s and 3 mm/s, and a post-test speed of 10 mm/s, sheer force of 5×10^−3^g and 5.0 mm distance, with the force of rupture or breaking (hardness) being recorded.

### Physicochemical Analysis

#### Proteins, lipids, total sugars, fibers, ashes and moisture

The centesimal composition was determined according to the methodology described by the Association of Official Analytical Chemist Methods [15]. The following tests were carried out: the micro-Kjedahl method, total sugars by the Fehling reduction, the fibers by Henneberg and Stohmann; moisture and total dry extract (EST) by drying in an oven stabilized at 105° C until obtaining constant weight; determination of ash content by carbonization followed by incineration in a muffle furnace at 550° C; determination of lipids by Folch, Lees, and Stanley (1957); and water activity was performed at 25° C in Aqualab® [16].

### Sensory evaluation

The sensorial tests were performed after approval by the Research Ethics Committee of the Health Sciences Center of the Federal University of Paraiba, under reference number 052900/2015. For Sensory Analysis, the Acceptance, Intention of Purchase and Ordering tests were performed by 123 untrained judges, being 76.4 2 % women and 23.57 % men. The samples were coded with random 3-digit numbers and served in plastic containers in a comparative way, in complete and balanced blocks. In addition to the samples, each evaluator received a glass with water to clean their palate during the evaluation. The tests were conducted at the Mauricio de Nassau Faculty (Joao Pessoa) and the Federal University of Paraiba, Joao Pessoa campus, both in the Dietetic Technique laboratory of each institution. The judges were between the ages of 18 and 61 and were in perfect health.

The judges received a sheet through which they evaluated each sample according to the attributes of appearance, color, aroma, flavor, texture and overall evaluation using a hedonic scale of 9 points, where 9 would be the maximum score represented by “I liked it very much”, and 1 is the minimum grade, representing “very disgusting”, containing intermediate points between these values. For purchase intent, participants responded to a 5-point scale expressing their willingness to consume, to buy or not buy the product offered to them, where terms were defined between “likely to buy” with a score of 5, and “probably would not buy” with a score of 1, and “maybe I would buy” at the intermediate point with a score of 3.

For the Ordination test carried out together with the Acceptance and Purchase Intention test, the judges were instructed to place the samples in ascending order according to their preference, with 1^st^ place being the most preferred and 4^th^ place the least preferred. Sensory tests were performed according to Dutcosky (2013) [17].

### Lipid profile of cookies

The samples of the cookies elaborated with different lipid sources (CVF - vegetable fat; CB - butter; CG - goat cream; CGCLA - goat cream with CLA) were analyzed for their lipid profiles. The transesterification procedure of the samples was carried out according to Sukhija and Palmquist (1988) and Palmquist and Jenkins (2003), with adaptations [12. 13]. The fatty acids were analyzed according to methyl esters by gas chromatography with flame ionization detection (GC-FID) using a QP2010-plus Shimadzu chromatograph (Shimadzu, Kyoto, Japan) equipped with a silica-fused (SP-2560, 100 m × 0.25 mm × 0.20 μm, Supelco, Bellefonte, PA, USA). During the analysis the injector and detector were maintained at 250°C and 280°C, respectively. Helium was used as the entrainment gas at a constant flow rate of 1 ml/min and 1 μl of sample was injected. The oven temperature was programmed to start at 50°C, then increased by 50°C every minute until it reached 150°C, remaining for 20 minutes, and next increased by 1°C every minute until it reached 190°C and remained for 1 minute, and then increased 2°C every minute until reaching 220°C, and finally remaining at 220°C for 30 minutes. The identified fatty acids were express as percentages and are shown in Table 1.

### Statistical analyses

The analysis data were submitted to analysis of variance (ANOVA) followed by Tukey test to verify if there was a significant difference between the samples, considering p <0.05 and using Sigma Stat 3.1 software [18]. The results of the sensory order-preference tests were analyzed according to the Friedman test using the Newell MacFarlane Table [19]. Physical and physico-chemical analyzes were performed in triplicate. All data sets were submitted to Principal Component Analysis (PCA) using correlation matrix and Statistica 7.0 software (Statsoft Inc., Tulsa, OK, USA).

## Results and discussion

### Physical analyzes

#### Color

The color data are shown in Table 3. Values of L* above 50 tend to be lighter in color as observed herein. Only the CB presented a difference in relation to the other formulations, with the lowest value (55.66), while the other treatments ranged from 58.34 (CGCLA), to 57.68 (CG) and 57.66 (CVF). Extreme values of L* may interfere with sensory perception, since they are either too dark or too light. The increase in the parameter a* and decrease in L* indicate the progress of the darkening, however this darkening associated to the increase of * and decrease of L* was not observed in cookies [20].

**Table 3.** Physical parameters of cookies with different lipid sources.

| Variable | Cookies |  |  |  |
| --- | --- | --- | --- | --- |
|  | CVF | CB | CG | CGCLA |
| **L** | 57,66<sup>a</sup>±0,87 | 55,66<sup>b</sup>±0,66 | 57,68<sup>a</sup>±0,99 | 58,34<sup>a</sup>±0,91 |
| **a\*** | 14,05 ±0,76 | 14,44 ±0,54 | 14,29 ±0,84 | 14,21 ±0,95 |
| **b\*** | 29,47<sup>c</sup>±0,91 | 30,57<sup>ab</sup>±0,99 | 29,76<sup>b</sup>±0,95 | 30,89<sup>a</sup>±0,94 |
| **Texture (Kg)** | 2,21<sup>c</sup>±0,45 | 5,54<sup>a</sup>±0,66 | 4,41<sup>b</sup>±0,42 | 4,05<sup>b</sup>±0,48 |

Means ± standard deviation with different letters on the same line differed by Tukey's test (p <0.05).
CVF - vegetable fat; CB - butter; CG - goat cream; CGCLA - goat cream with CLA.

The parameter a* presented no significant difference (p> 0.05), however it did obtain positive values which are associated with red pigments, since the parameter b* obtained a significant difference between the formulations; furthermore, the values were positive, being related to yellow pigments. According to Marques (2016), positive values of a* and b* are expected in cakes, since they present red and yellow pigments, respectively, due to the caramelization reactions of sugars and Maillard [21].

The Maillard reaction is an interaction between sugars and amino acids, and together with the caramelization reaction of sugars produce brown pigments during the cooking process, which are associated with luminosity, and more red and yellow staining [22].

The variation in L* and b* did not interfere in the sensorial perception of the appearance and color (Table 5), as will be presented in the subitem “Consumer Test”. According to Sharma and Gurjral (2013), browning reactions are influenced by other factors such as moisture, water activity, pH, sugars, type and proportion of starch, cooking time and temperature [23].

**Table 4.** Physico-chemical parameters of cookies made with different lipid sources.

| Variable | Cookies |  |  |  |
| --- | --- | --- | --- | --- |
|  | CVF | CB | CG | CGCLA |
| **Lipids** | 28,00<sup>a</sup> ± 0,00 | 25,80<sup>b</sup> ± 0,66 | 21,16<sup>c</sup> ± 0,24 | 20,86<sup>c</sup> ± 0,68 |
| **Proteins** | 5,43 ± 0,25 | 5,50 ± 0,32 | 5,90 ± 0,08 | 5,79 ± 0,30 |
| **Total Sugars** | 51,38<sup>c</sup> ± 0,32 | 54,42<sup>b</sup> ± 0,36 | 56,38<sup>a</sup> ± 0,33 | 55,02<sup>b</sup> ± 0,12 |
| **Fibers** | 0,24 ± 0,01 | 0,25 ± 0,01 | 0,26 ± 0,01 | 0,25 ± 0,00 |
| **Ashes** | 1,61 ± 0,24 | 1,70 ± 0,16 | 1,63 ± 0,27 | 1,59 ± 0,28 |
| **Humidity** | 4,39<sup>c</sup> ± 0,29 | 4,64<sup>c</sup> ± 0,22 | 6,52<sup>a</sup> ± 0,23 | 5,35<sup>b</sup> ± 0,07 |
| **Aw** | 0,35<sup>b</sup> ± 0,47 | 0,43<sup>ab</sup> ± 0,03 | 0,53<sup>a</sup> ± 0,17 | 0,47<sup>ab</sup> ± 0,26 |

Averages ± standard deviation with different letters on the same line differed by Tukey's test (p<0.05). CVF - vegetable fat; CB - butter; CG - goat cream; CGCLA - goat cream with CLA.

**Table 5.** Mean values of sensory acceptance and purchase intention tests.

| Attribute | Cookies |  |  |  |
| --- | --- | --- | --- | --- |
|  | CVF | CB | CG | CGCLA |
| Appearance | 7,47 ±1,36 | 7,35 ±1,43 | 7,46 ±1,45 | 7,32 ±1,38 |
| Color | 7,62 ±1,18 | 7,37 ±1,32 | 7,41 ±1,40 | 7,40 ±1,29 |
| Aroma | 6,83 ±1,86 | 6,83 ±1,70 | 7,09 ±1,51 | 6,72 ±1,90 |
| Flavor | 6,81 ±2,12 | 6,80 ±1,83 | 7,15 ±1,67 | 6,96 ±1,76 |
| Texture | 7,00 ±1,98 | 7,03 ±1,58 | 7,14 ±1,75 | 7,03 ±1,56 |
| Overall assessment | 7,20 ±1,62 | 7,13 ±1,48 | 7,53 ±1,32 | 7,10 ±1,60 |
| Purchase intention | 3,60<sup>ab</sup> ±1,30 | 3,52<sup>b</sup> ±1,22 | 3,96<sup>a</sup> ±1,11 | 3,67<sup>ab</sup> ±1,26 |

Averages ± standard deviation with different letters on the same line differed by Tukey's test (p<0.05). CVF - vegetable fat; CB - butter; CG - goat cream; CGCLA - goat cream with CLA.

### Instrumental texture

The cookies’ hardness (evaluated by the instrumental method) is proportional to the force applied to cause deformation. This strength varies according to the composition (quality and quantity of flour, sugars, fats, liquids and other ingredients), cooking time and temperature, humidity, and storage conditions. Hardness is one of the factors that is associated with texture to determine the acceptability of foods, so it is desirable that their values are not very high [24]. The values of the instrumental texture are presented in Table 3.

The CB formulation presented the highest hardness value (5.54 kg) and CFV presented the lowest value (2.21 kg). The CG and CGCLA formulations presented values of 4.41 kg and 4.05 kg, respectively (p> 0.05). These data demonstrate that the cookies made with the goat cream independent of the CLA content did not differ statistically from each other. An extreme increase of hardness as very soft or very hard can sensorially interfere in the products. Fragile and crunchy foods are known to have irregular and irreproducible relationships with the force versus deformation relationship [25].

The results obtained in the present study were similar to that of the control group (C). The results showed that the order of hardness was MGCO>C>GCO, and with the addition of oil (GCO) or microencapsulated oil (MGCO), the acceptance was C>MGCO>GCO, with the most accepted sample being that which obtained an intermediate degree of hardness [26]. According to Bertolin (2013), high values in texture negatively influence the overall acceptance of cookies. In the present work, this relationship of hardness with acceptance was not observed, considering that there was no significant difference in the sensory perception of texture (Table 5) [27]. Regarding hardness, the order was CB>CG = CGCLA>CFV (Table 3).

### Physico-chemical analysis

The physicochemical data are shown in Table 4, in which CVF and CB presented the highest lipid percentages with 28 % and 25.8 %, respectively. The fact that CVF and CB have higher lipid levels is related to the lower percentages of sugars and moisture. A correlation of the fat processing with the lipid content variation of the cookies can be made, since the cream is the emulsion of fat in water and the lipid content can vary from 12 to 60 %, since other elements like sugars can pass next to the fat globules in the milk skimming. Unlike the cream, the butter should contain a minimum percentage of 80 % lipids, and the hydrogenated vegetable fat is composed of 100 % lipids [28].

The humidity and Aw presented significant differences, with CG and CGCLA having the highest humidity values with 6.25 % and 5.35%, respectively, and CG had the highest Aw (0.53). These higher percentages can be associated with the characteristic of the lipids used in these formulations (Table 1), as previously mentioned. These variations may occur because other elements such as diversity and type of ingredients, cooking time and temperature directly interfere with moisture and Aw. Protein, fiber and ash contents did not vary significantly between formulations.

In the case of oatmeal-rich biscuits, the water content of the oatmeal oil was similar to that of oysters. According to Chung, Cho, and Lim (2014), the moisture of the biscuit cookie should vary between 2 and 8 g/100 g [29]. Therefore, the four cookie formulations are suitable for this variation. Foods have a sigmoid relationship with water and Aw, since the moisture content and Aw have a strong effect on the perception of sharpness and the mechanical sensations of dry and brittle foods as in biscuits [30, 31].

### Sensory evaluation

The evaluation of the mean scores obtained in the sensory analysis (Table 5) showed that the formulations did not differ significantly between the parameters (appearance, color, aroma, flavor, texture and overall evaluation) (p> 0.05). Despite the use of different types of fats with different and peculiar characteristics such as goat cream in the preparation of cookies, these changes were not perceptibly noticeable to the judges. The formulations were only qualitatively and not quantitatively altered in relation to the fat added to the cookies (Table 1), although there was a significant difference (p <0.05) in the lipid content of these formulations (Table 4).

When developing biscuits formulated with different concentrations of oats and palm oil, Bertolin et al. (2013) found that higher concentrations of oats (51 %) and lower concentrations of palm oil (8 %) negatively interfered in the sensorial acceptance of this biscuit [27]. Similar data regarding the fat content in biscuits were observed by Biguzzi, Schlich and Lange (2014) when evaluating the sensorial perception in biscuits with a reduction of different concentrations of fats (15 % and 25 %) compared to the standard biscuits, observing that the cookies made with lower fat percentage (25 % reduction) obtained sensorial difference in relation to the formulation with lower fat reduction and the standard formulation [32].

The difference in sensory perception in the reduction of the fat content in cookies was observed by Drewnowski, Nordensten and Dwyer (1998), who reported the decrease in the acceptance of biscuits with a reduction of 50 % fat; a fact that did not interfere when the reduction was 25 % [33].

The references mentioned above demonstrate that the amount of fat above a reduction percentage interferes in the judges’ acceptance. However, through this study it was observed that the variation in fat type could not be perceived by the judges, as there was no interference in the sensorial acceptance, or a significant difference among the evaluated attributes. Thus, the use of fats with a better lipid profile such as G and GCLA (Table 1) offers great nutritional and technological impact, adding value to the cookies without altering their flavor.

Among the analyzed attributes, the biscuit elaborated with the goat cream presented greater intention of purchase, resembling the cookies containing vegetable fat and goat cream enriched with CLA (p<0.05). In addition, all formulations scored between 4 “I would possibly buy” and 3 “I would maybe buy/I would maybe not buy,” with at least a 50 % chance of acquiring cookies.

According to the Prediction Order Analysis data (Table 6), there was no significant difference between the four formulations (p> .05). The CG and CGCLA samples are as preferred as the CVF, which was elaborated with VF, and which is more used in the preparation of biscuits and cookies by industries [34]. These data indicate that cookies can be elaborated from better sources of fat such as goat cream, which has a better lipid profile (Table 1) and which brings benefits to consumer health. In addition, the characteristic odors and flavors of goat origin products were not sensorially perceived when compared to the other formulations.

**Table 6.** Distribution of the scores according to the order of general preference by the tasters (n = 123) in the sensory analysis of cookies.

| Cookies formulations | Number of testers by order* |  |  |  | Total orders** |
| --- | --- | --- | --- | --- | --- |
|  | 1 | 2 | 3 | 4 |  |
| CVF | 32 | 27 | 22 | 42 | 320 |
| CB | 34 | 36 | 34 | 19 | 284 |
| CG | 22 | 35 | 27 | 39 | 329 |
| CGCLA | 35 | 25 | 40 | 23 | 297 |

\* 1 = least preferred, 4 = most preferred.
\*\* Sum of the orders of each sample = (1 x number of tasters) + (2 x number of tasters) + (3 x number of tasters) + (4 x number of tasters).
CGV - Vegetable fat; CM - Butter; Caprine CNCLA - Goat cream enriched with CLA).

The use of goat cream is a lipid source that can be applied to cookies, thus adding value and functionality to goat milk derivatives, which are normally used in dairy products. In this way, the present work will serve as a subsidy for the use of this fat in developing various confectionery and bakery products.

### Chromatographic analysis of cookies

The fatty acid profiles of the cookies are shown in Table 7. The CB, CG and CGCLA presented higher percentages of SFA in comparison to the CVF. However, when comparing CG with CGCLA, it is observed that CGCLA reduced 7.89 % of the SFA, and this reduction is associated with the addition of soybean oil in the goat diet, which altered the lipid profile of goat cream [11].

**Table 7.**
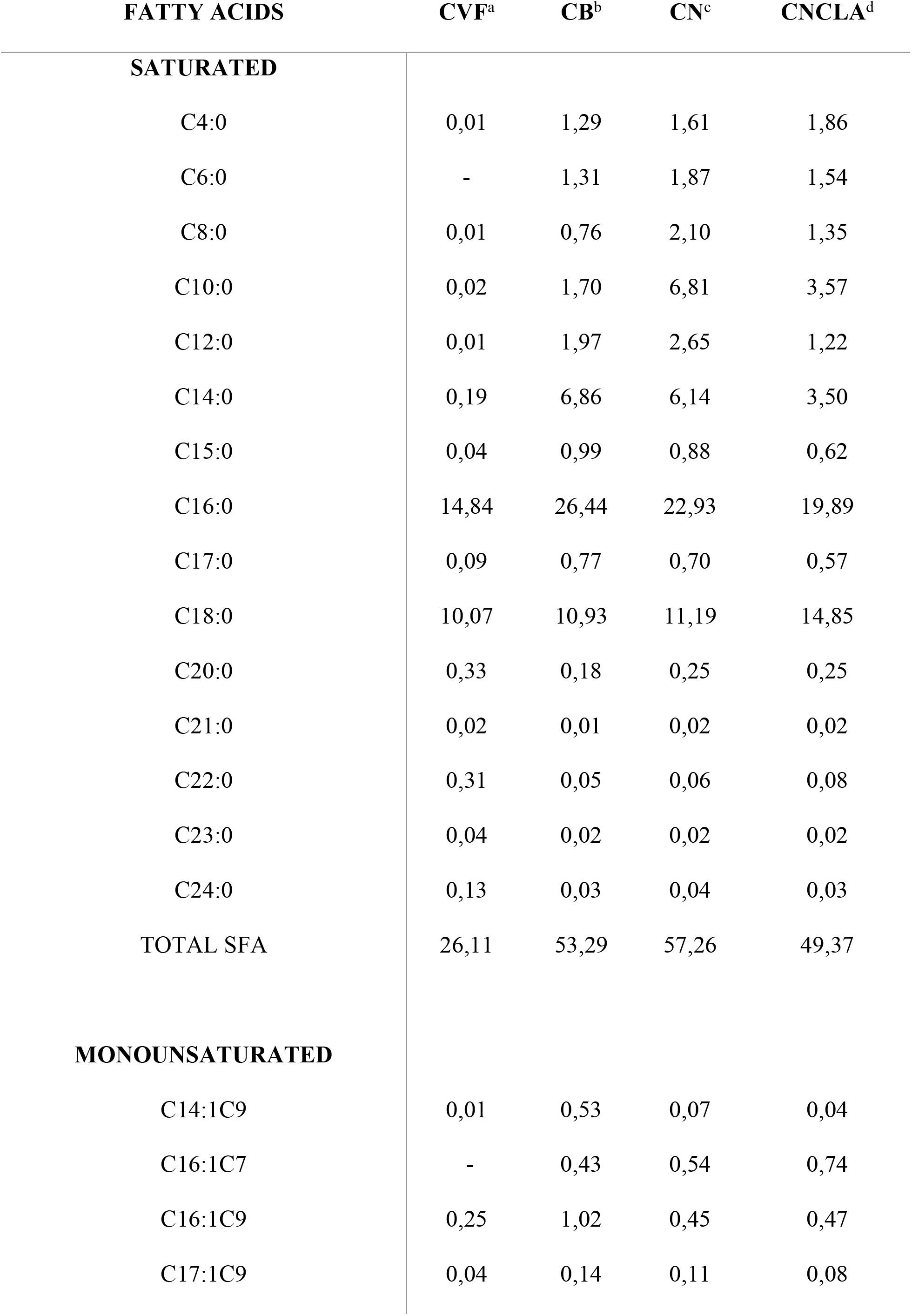
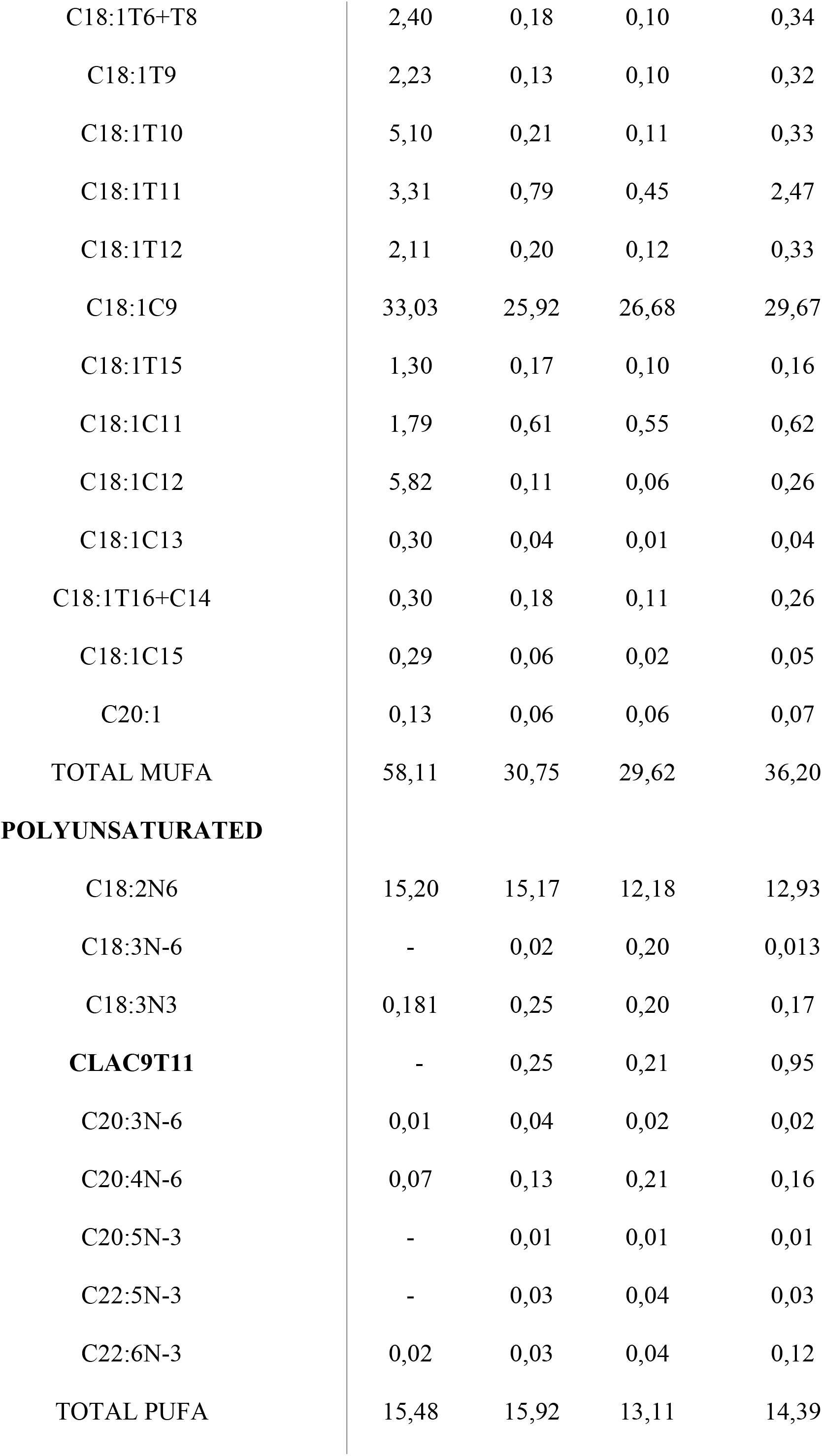
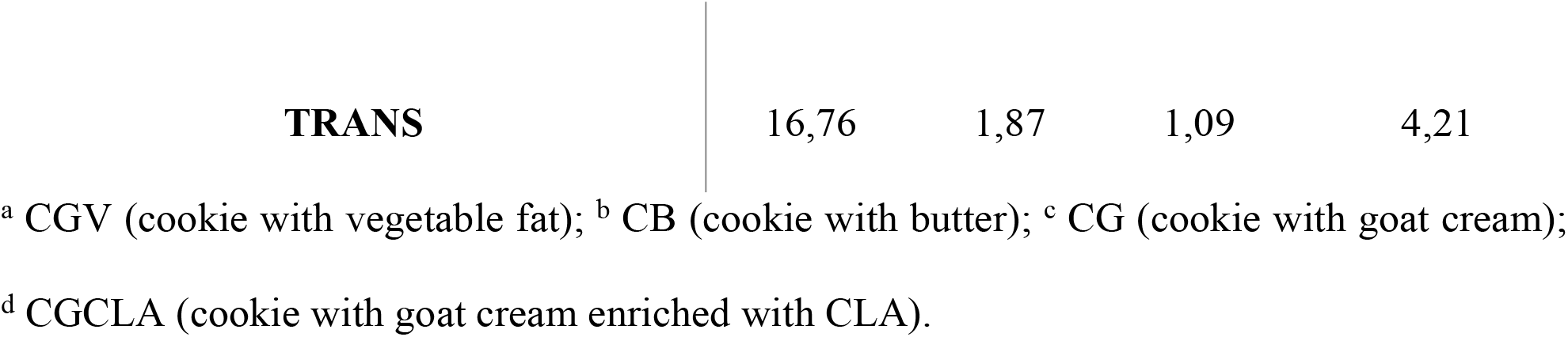
Profile of fatty acids present in cookies in percentage.

It is common for fats of animal origin such as those used in these cookies to be a source of SFA. Palmitic acid was the predominant FA in CB, CB and CGCLA, representing about 20 % of total FA. Capric acid (C10: 0) and caprylic (C8: 0) acids are characteristic of goat fat, being identified in greater proportions in G with 11.14 % and 3.27 %, and in GCLA with 5.28 % and 1.84 %, respectively (Table 1).

The decrease of the medium chain fatty acids (MCFA) (10 to 16 carbons) is associated with the manipulation of the goat diets, since the addition of calcium salts of fatty acids in the goat’s feed can present a decrease in the capric and myristic concentration and one an increase in linoleic fatty acids (C18: 2 n6), CLA (C18: 2 c9t11), omega-6 (n-6) and PUFA [35]. In the elaborated cookies in the present research, the decrease of C10: 0 and the increase of CLA were also perceived in both CG and CGCLA. This fat naturally enriched with CLA used in the elaboration of the CGCLA also obtained similar treatment, where the caprine ration was added with soybean oil [11].

High consumption of SFA is commonly associated with risk factors for cardiovascular disease. However, they should not be excluded from feeding, since they are necessary for energy synthesis and make up the cell membranes.^36^ Short chain saturated fatty acids (4 to 8 carbons) also exert beneficial effects on intestinal health due to rapid uptake and oxidation by colon cells, thereby stimulating cell proliferation in this tissue, as well as playing a role in disease prevention for cardiovascular, cancer and inflammatory diseases [37].

The MUFA presented a higher percentage in CGV with 58.11 %, and these values were related to the TFA, which obtained high percentages when compared to CB, CG and CGCLA. The iTFA present in vegetable fat are associated with CNCD, since the organism does not have the capacity to metabolize these compounds, leading to atheromas. The TFA content ranged from 16.76 % (CVF) to 1.1 % (CG), with CGCLA being the second highest percentage of TFA with 4.21 %. Although CGCLA showed an increase in TFA when compared to CB and CG, this change in lipid profile increased in the cookies related to the fat used in its formulation, which has a higher percentage of essential fatty acids such as CLA.

The high percentage of AGV of CGV is related to the lipid profile of GV used in the elaboration of CGV. The process of vegetable fat hydrogenation increases the percentage of industrial trans fatty acids (iTFA), with these fatty acids being found in industrialized foods such as cookies and wafers that use this fat in their composition.^34^ When evaluating the percentages of TFA in processed foods in the Portuguese market, Costa et al. (2016) observed that Trans-C18: 1 and its isomers were present in higher amounts in these foods.^7^

Elaidic acid (C18: 1 trans-9) is the main constituent of iTFA. Oleic acid (C18: 1) is an MUFA present in oils of vegetable origin. After the hydrogenation process, the oleic acid changes its spatial and biochemical conformation, forming elaidic acid.^38^ These iTFA are associated with cardiovascular diseases, obesity, and CNCD. ^39^

The PUFA percentage did not have much quantitative variation, with the highest percentage of CVF (15.48 %) and the lowest of CG (13.10 %); however, the lipid profile qualitatively differs between formulations. Among the PUFA, CGCLA and CG presented higher percentages of docosahexaenoic acid (DHA) (C22: 6n-3), which is associated with cardio-protective effect, neurological development, and anti-inflammatory properties.^26^

Another essential PUFA is CLA (C18: 2 c9t11), which was only present in ruminant fat, since isomerization and biohydrogenation of linoleic acid (C18: 2) and linolenic acid (C18: 3) only occur in rumen or mammary glands of these animals. Meat or dairy products are the main source of consumption of these TFAr [40, 41].

This natural transfat derived from ruminants exerts functional health properties such as plasma antioxidant activity, antimutagenic and anticarcinogenic action, acting in the reduction of cytotoxic agents in cancer cells. In addition, CLA triggers immune response stimuli against atherosclerosis, diabetes mellitus and obesity prevention [3, 42].

After the preparation of the cookies, there was a reduction in CLA contents of 74 % compared to fat enriched with CLA before cooking. This loss occurred in all cookies containing CLA, however, the CGCLA still maintained a higher content of 70 % of this fatty acid compared to the others. Therefore, the cookie with caprine enriched with CLA is an alternative for the consumption of this essential fatty acid, since CLA is usually consumed through dairy and meat products. Thus, the cookies with CLA diversify and add value to products of goat origin and can help to produce a food with functional potential.

### Principal component analysis

Figure 1 shows the graph resulting from Principal Component Analysis (PCA). The first two factors explain 90.03 % of the variance, allowing relevant discrimination of the samples as a function of the attributes evaluated, according to Piclin et al. (2008) and Nurgel et al. (2004) [43, 44]. The first factor (A) explained 56.29 % of the variance, being positively characterized by “b*”, “proteins”, “sugars”, “moisture”, “ashes”, “fibers”, “Aw”, “purchases”, “SFA”, “overall assessment” and “purchase intention”, and negatively by “lipids”, “MUFA” and “TFA”. The second factor (B) explained 33.74 % of the total variance, relating positively to “L”, and negatively by “texture”, “ashes” and “PUFA”.

**Fig 1.**
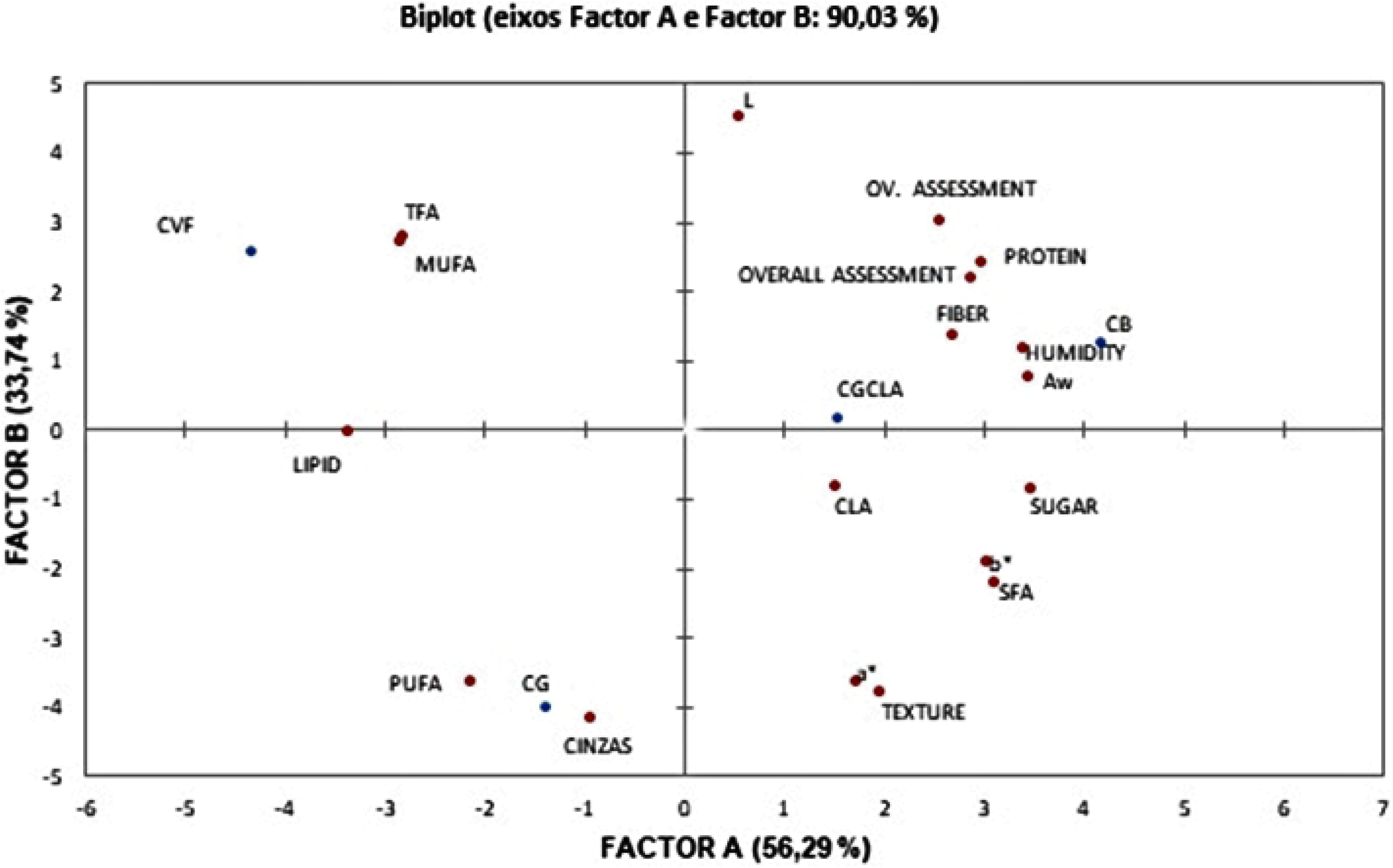
PCA of the data for physical-chemical, physical, sensory and chromatographic of the cookies. CVF (cookie with vegetable fat); CB (cookie with butter); CG (cookie with goat cream) and CGCLA (cookie with enriched goat cream with CLA).

It can be observed that the analysis was able to separate the samples according to similar characteristics, with the CVF formulation correlated with the monounsaturated fatty acids and trans fatty acids; CB was related to polyunsaturated fatty acids and ashes; the CG and CGCLA formulations obtained more attributes in common such as Aw, moisture, proteins, fibers, purchase intention and overall evaluation. CGCLA was correlated with CLA, considering that this formulation was the one that obtained the highest percentage of this PUFA. The principal components among the four formulations corroborate the data obtained in the analyzes described above.

## Conclusion

The substitution of the hydrogenated vegetable fat and butter by goat cream in cookies is feasible, since it does not alter the flavor, maintaining the sensorial acceptance and improving the profile of fatty acids of the food, and increasing the levels of essential fatty acids like CLA. This fact raises the biological properties of this food and may increase the demand for this product by consumers seeking to eat tasty foods with functional potential.

## Acknowledgments

This study was supported by Coordination of Improvement of Higher Level Personnel - CAPES.

## Supporting information

**S1 Table. Fatty acid profile present in fats in percentage.** ^a^ VF: vegetable fat; ^b^ B: butter, ^c^G: goat cream; ^d^ GCLA: goat cream with CLA, expressed as a percentage.

**S2 Table. Ingredients used in the formulations of cookies for 100 g of fresh dough.** ^a^ CVF-vegetable fat; ^b^ CB - butter; ^c^ CG - goat cream; ^d^ CGCLA - goat cream with CLA.

**S3 Table. Physical parameters of cookies with different lipid sources.** Means ± standard deviation with different letters on the same line differed by Tukey’s test (p <0.05). CVF - vegetable fat; CB - butter; CG - goat cream; CGCLA - goat cream with CLA.

**S4 Table. Physico-chemical parameters of cookies made with different lipid sources.** Averages ± standard deviation with different letters on the same line differed by Tukey’s test (p<0.05). CVF vegetable fat; CB - butter; CG - goat cream; CGCLA - goat cream with CLA.

**S5 Table. Mean values of sensory acceptance and purchase intention tests.** Averages ± standard deviation with different letters on the same line differed by Tukey’s test (p<0.05). CVF vegetable fat; CB - butter; CG - goat cream; CGCLA - goat cream with CLA.

**S6 Table. Distribution of the scores according to the order of general preference by the tasters (n = 123) in the sensory analysis of cookies.** * 1 = least preferred, 4 = most preferred.

** Sum of the orders of each sample = (1 × number of tasters) + (2 × number of tasters) + (3 × number of tasters) + (4 x number of tasters).

CGV - Vegetable fat; CM - Butter; Caprine CNCLA - Goat cream enriched with CLA).

**S7 Table. Profile of fatty acids present in cookies in percentage.** ^a^ CGV (cookie with vegetable fat); ^b^ CB (cookie with butter); ^c^ CG (cookie with goat cream); ^d^ CGCLA (cookie with goat cream enriched with CLA).

**S1 Fig. PCA of the data for physical-chemical, physical, sensory and chromatographic of the cookies**. CVF (cookie with vegetable fat), CB (cookie with butter), CG (cookie with goat cream) and CGCLA (cookie with enriched goat cream with CLA).

